# Host immune responses to enzootic and invasive pathogen lineages vary in magnitude, timing, and efficacy

**DOI:** 10.1101/2021.12.08.471676

**Authors:** Coby A. McDonald, C. Guilherme Becker, Carolina Lambertini, L. Felipe Toledo, Célio F.B. Haddad, Kelly R. Zamudio

**Author notes:** Department of Microbiology, Immunology, and Pathology, Colorado State University, Fort Collins, CO 80523, USA.

## Abstract

Infectious diseases of wildlife continue to pose a threat to biodiversity worldwide, yet pathogens are far from monolithic in virulence. Within the same pathogen species, virulence can vary considerably depending on strain or lineage, in turn eliciting variable host responses. One pathogen that has caused extensive biodiversity loss is the amphibian-killing fungus, *Batrachochytrium dendrobatidis* (Bd), which is comprised of a globally widespread hypervirulent lineage (Bd-GPL), and multiple geographically restricted lineages. Whereas host immunogenomic responses to Bd-GPL have been characterized in a number of amphibian species, immunogenomic responses to geographically-restricted, enzootic Bd lineages are unknown. To examine lineage- specific host immune responses to Bd, we exposed a species of pumpkin toadlet, *Brachycephalus pitanga*, which is endemic to Brazil’s Southern Atlantic Forest, to either the Bd-GPL or the enzootic Bd-Asia-2/Brazil (hereafter Bd-Brazil) lineage. We quantified functional immunogenomic responses over the course of infection using differential gene expression tests and coexpression network analyses. Host immune responses varied significantly with Bd lineage. Toadlet responses to Bd-Brazil were weak at early infection (26 genes differentially expressed), peaked by mid-stage infection (435 genes) and were nearly fully resolved by late-stage disease (9 genes). In contrast, responses to Bd-GPL were magnified and delayed; toadlets differentially expressed 97 genes early, 86 genes at mid-stage infection, and 728 genes by late-stage infection. Given that infection intensity did not vary between mid- and late-stage disease, this suggests that pumpkin toadlets may be at least partially tolerant to the geographically-restricted Bd- Brazil lineage. In contrast, mortality was higher in Bd-GPL-infected toadlets, suggesting that late-stage immune activation against Bd-GPL was not protective and was consistent with immune dysregulation previously observed in other species. Our results demonstrate that both the timing of immune response and the particular immune pathways activated are specific to Bd lineage. Within regions where multiple Bd lineages co-occur, and given continued global Bd movement, these differential host responses may influence not only individual disease outcome, but transmission dynamics at the population and community levels.

## INTRODUCTION

Host responses to infection are important in determining individual disease outcomes, which have consequences for both host and pathogen fitness, transmission dynamics, and coevolution (Råberg et al., 2009a). Likewise, the immunological mechanisms conferring host tolerance, resistance, and susceptibility can vary depending on host and pathogen genotype, as well as biotic and abiotic environmental factors (Dobson, 2004; Hawley & Altizer, 2011). Host-pathogen interactions are further complicated in pandemic scenarios, in which novel pathogen genotypes evolve and gain access to naïve hosts. Thus, differentiating individual host responses to enzootic and invasive pathogen lineages is important for understanding host-pathogen interactions at the population and community scale.

Host tolerance, resistance, and susceptibility are especially complicated in the amphibian-*Batrachochytrium dendrobatidis* (Bd) system. Chytridiomycosis, the disease caused by Bd, has triggered panzootic declines in more than 500 amphibian species since the 1970s (Scheele et al., 2019). Such catastrophic population declines are thought to be linked to human-mediated spread of a hypervirulent lineage, Bd-GPL, within the 20^th^ century (O’Hanlon et al., 2018). Whereas Bd-GPL has caused epizootics on all continents harboring amphibians, five less virulent, enzootic lineages are also present in East Asia (Bd-Asia1), Europe (Bd-CH), South Africa (Bd-Cape), Brazil (Bd- Asia2/Brazil), and Korea (Bd-Asia3) (Byrne et al., 2019; O’Hanlon et al., 2018; Scheele et al., 2019). Native host persistence in regions populated by enzootic Bd is thought to be due to species-specific defense strategies ranging from resistance to tolerance (Bataille et al., 2013; Greenspan et al., 2018; O’Hanlon et al., 2018).

Among geographic locations where enzootic Bd lineages persist, host disease trajectories and pathogen dynamics have been comprehensively studied in Brazil’s Atlantic Forest (Jenkinson et al., 2016; Rodriguez et al., 2014). Although less than 16% of its original extent remains, the Atlantic Forest harbors nearly 10% of all amphibian species and high levels of amphibian endemism (Haddad et al., 2013; Toledo et al., 2014). Population and species-level declines in the region have been linked to Bd, the pathogen has been present since at least the late 1800s, and it is one of few locations where enzootic and Bd-GPL lineages are known to co-occur and hybridize (Byrne et al., 2019; Carvalho et al., 2017; Jenkinson et al., 2016; Rodriguez et al., 2014; Schloegel et al., 2012).

Host responses to Bd are species-specific and vary based on life history, range, and pathogen genotype. When infected with virulent Bd-GPL, direct developing, native Brazilian hosts experience higher infection intensity and mortality than native species with aquatic life stages (Mesquita et al., 2017). When exposed to a less virulent Bd-GPL genotype, native species exhibit no mortality, but non-lethal disruptions in water balance (Bovo et al., 2016). Host survival is significantly higher in both direct-developing and aquatic breeding Brazilian hosts exposed to Bd-Brazil relative to Bd-GPL or hybrid genotypes (Greenspan et al., 2018). Notably, while enzootic Bd-Brazil causes little mortality in Brazilian hosts, it causes higher mortality rates in non-native species (Becker et al., 2017; Greenspan et al., 2018). Enzootic Bd-Brazil genotypes are currently being outcompeted by invasive Bd-GPL, which may rapidly shift community dynamics in the region (Jenkinson et al., 2016; Jenkinson et al., 2018).

While functional responses to enzootic Bd have not been explicitly tested, host susceptibility and resistance to Bd-GPL has been characterized in several species (Grogan, Robert, et al., 2018; Zamudio et al., 2020). Host resistance and consequent survival is mediated both by timing and magnitude of immune responses. Multiple studies have found that susceptible hosts upregulate more immune genes to a greater magnitude than do resistant hosts (Ellison, Tunstall, et al., 2014; Eskew et al., 2018; Poorten & Rosenblum, 2016). When testing for timing of response, Grogan, Cashins, et al. (2018) found that more resistant frog populations increased expression of both innate and adaptive immune genes earlier during the period that infection was still subclinical, whereas susceptible populations mounted strong immune responses at late- stage infection. While susceptible species do mount robust responses to infection that may include epithelial remodeling at the site of infection and cytokine, complement, cell- mediated, and antibody-mediated responses both in the skin and systemically, these late responses are insufficient to decrease pathogen load and resolve clinical disease (Ellison, Savage, et al., 2014; Eskew et al., 2018; Grogan, Cashins, et al., 2018).

Given previous work outlining functional mechanisms of resistance versus susceptibility in various host species infected with Bd-GPL, we sought to further differentiate between host responses to enzootic Bd-Brazil or invasive Bd-GPL by examining both temporal and lineage-specific responses. We measured transcriptome- wide responses in immunologically-relevant tissues (skin, spleen, and intestine) at three time points over the course of infection to identify 1) shared host responses irrespective of Bd lineage, 2) lineage-specific immune strategies, and 3) variation in timing of immune response.

## MATERIALS AND METHODS

### Study species

We examined host Bd responses of the pumpkin toadlet, *Brachycephalus pitanga* (Brachycephalidae), a direct-developing frog that is endemic to the Southern Atlantic Forest of Brazil. Because the pumpkin toadlet does not have an aquatic life stage, environmental exposure to the waterborne Bd fungus is thought to be low. However, this species occurs at very high population densities within its restricted range in and near Parque Estadual da Serra do Mar, São Luiz do Paraitinga, state of São Paulo, Brazil, potentially increasing probability of direct transmission. Pumpkin toadlet susceptibility to Bd appears to be strain-specific; one study demonstrated high infection load, susceptibility, and mortality upon exposure to Bd-GPL strain CLFT159, whereas another reported no significant mortality or sub-lethal effects with Bd-GPL strain CLFT023 (Bovo et al., 2016; Mesquita et al., 2017).

### Infection trial

Adult *B. pitanga* were collected from private property in São Luiz do Paraitinga, state of São Paulo, and acclimated to laboratory conditions for two weeks prior to infection trial (Cornell University IACUC #2014-0042, CEUA-Unicamp # 5796-1, SISBIO # 17242).

Following acclimation, frogs were swabbed for preexisting Bd infection; all animals were confirmed Bd-negative via qPCR. Frogs were housed individually in plastic terraria at a constant temperature of 19°C with a 12:12 light:dark cycle. Each terrarium contained autoclaved sphagnum moss kept moist with deionized and aged water. Frogs were fed pinhead crickets and locally-collected small arthropods *ad libitum* throughout the trial.

As part of a larger study, we randomly assigned 78 frogs to one infection treatment: invasive Bd (Bd-GPL; CLFT159), enzootic Bd (Bd-Brazil; CLFT001), and controls. *Bd*-exposed frogs were individually inoculated with 3.9x10^6^ zoospores of either Bd-GPL or Bd-Brazil, which was the maximum number of zoospores per frog that we could standardize between Bd strains. Inoculum was suspended in 5 mL of 1% tryptone broth and frogs were exposed for 30 minutes in petri dishes that ensured ventral skin contact. Control frogs were sham-inoculated via exposure to 1% tryptone broth for 30 minutes. We swabbed frogs every five days to monitor infection load over the course of the experiment. We quantified load via qPCR according to Hyatt et al. (2007). This protocol targets the Internal Transcribed Spacer (ITS) region of Bd. Because ITS copy number varies with Bd genotype and can confound quantification, we used a gBlock® gene fragment to generate standard curves and estimate ITS copies present (Hyatt et al., 2007; Longo et al., 2013).

*Brachycephalus pitanga* has a short course of chytridiomycosis infection, with a recent study demonstrating 100% mortality within 10 days (Mesquita et al., 2017). To capture variation in immune response over the course of Bd infection, we euthanized frogs from each treatment at 5, 10, and 15 days post-infection. We harvested skin, liver, and intestine within three minutes of euthanasia and stored tissue samples in RNAlater at -80°C for RNA isolation. Because *B. pitanga* has a snout-to-vent length of less than 2 cm, we were unable to isolate splenic tissue, which is the major lymphoid organ in adult frogs. To best capture the immune repertoire, we instead measured expression differences in skin, which is the site of infection, as well as liver and intestine, the latter of which has minor lymphoid function in the gut-associated lymphoid tissue (GALT).

Because preexisting natural infection could confound gene expression responses, we included only frogs that were Bd-free at the start of the trial for RNA-sequencing. This resulted in a final sequencing dataset of 108 tissue samples such that we included a minimum of four biological replicates per treatment, time, and tissue combination.

### Library preparation and sequencing

We isolated RNA from all tissue samples using RNeasy Mini Kits (Qiagen, CA, USA) according to the manufacturer’s protocol and including a 15-minute on-column DNase digestion. We quantified RNA using a Qubit 2.0 (ThermoFisher Scientific, CA, USA), and confirmed RNA integrity via gel electrophoresis and Fragment Analyzer (RQN: 8.4- 10; Agilent Technologies, Inc, CA, USA). We standardized samples to 75 ng RNA for library preparation input. All libraries were prepared using NEBNext Poly(A) mRNA magnetic isolation modules and Ultra II Directional Library Prep kits (New England BioLabs, Inc, MA, USA) according to standard protocol. Libraries were amplified with 16 PCR cycles, and library quality and fragment size were confirmed on a Fragment Analyzer. We sequenced libraries on a NextSeq 500 (Illumina, CA, USA) over three lanes: one lane of paired-end, 2x75 bp sequencing for de novo transcriptome assembly, and three lanes of single-end, 75 bp sequencing for alignment and quantification.

### Brachycephalus pitanga transcriptome assembly and annotation

We assessed quality of raw reads using FastQC (v 0.11.8). Raw reads were then trimmed with Trim Galore (v0.6.3; https://github.com/FelixKrueger/TrimGalore) such that only reads ≥ 36 bp that met a quality score of 5, a stringency of 1, and an error rate of 0.1 were retained. Trimmed read quality was again assessed via FastQC. We assembled the *B. pitanga* transcriptome with Trinity (v 2.8.4), using only trimmed, paired-end liver, skin, and intestine reads and default Trinity parameters. Transcriptome quality was assessed with BUSCO (v 3.1.0) and Trinity-packaged scripts to compute Nx and ExN50 statistics.

The assembled raw transcriptome was annotated using Trinotate version 3.0.0. Briefly, we identified putative coding regions using TransDecoder version 5.4.0 with blastp and Pfam homology searches. We then performed blastx and blastp, Pfam homology, SignalP, tmHMM, and RNAMMER searches using either the transcriptome or predicted open reading frame files according to Trinotate specifications. Following annotation, we removed all transcripts with non-vertebrate annotations, retaining only unannotated and vertebrate-annotated transcripts as our final, cleaned *B. pitanga* transcriptome.

We aligned all single-end reads to our cleaned *B. pitanga* transcriptome using bowtie2 (v2.3.5.1) and quantified read abundance using RSEM (v1.2.29) in Trinity. We assessed for batch effects and sample outliers using PCA.

### Gene expression analyses

To identify tissue-specific differential responses to enzootic and epizootic *Bd* over time, we performed pairwise differential expression tests in edgeR (v 3.23.3) using gene-level raw read counts. After removing outliers, we partitioned samples according to tissue and analyzed each tissue separately. We normalized samples according to library size and retained only genes that were expressed at ≥ 1 counts per million (CPM; ≥ 8 absolute reads given minimum library size) in at least two-thirds of all samples.

Following read filtering, we estimated common and tagwise dispersion, fit generalized linear models, and performed likelihood ratio tests on all pairwise comparisons of interest. Pairwise comparisons were performed both within treatment over time (e.g. Bd- GPL^+^ early versus Bd-GPL^+^ late), between treatment and control at each time point (e.g. Bd-Brazil-mid versus Control-mid). Following initial comparisons, we additionally compared mid-stage Bd-Brazil^+^ and late-stage Bd-GPL^+^ frogs to dissociate temporal and pathogen-specific responses. Gene expression was considered significant using a false discovery rate (FDR) corrected *p* value of < 0.05.

We further analyzed expression responses using Weighted Gene Co-Expression Network Analysis (WGCNA), which is able to identify subtler, yet potentially equally biologically relevant gene expression changes than is possible with pairwise differential expression tests (Langfelder & Horvath, 2008). First, we reduced background signals in our gene expression dataset by filtering with variancePartition (v. 1.12.3).

VariancePartition estimates the percent of variation attributable to variables of interest using linear mixed models; genes with high residual variation can then be removed. For each tissue we removed batch effects by considering batch as a random effect in an initial LMM and retaining residuals. Retained residuals were then imputed in the second LMM with categorial variables (treatment and time) set as random effects and a continuous variable (infection load) set as a fixed effect. We retained all genes above the 90^th^ quantile of expression variation distribution for each trait. Filtered gene counts were then log2-transformed and used as input for WGCNA.

We constructed signed gene co-expression networks for each tissue separately using standard WGCNA stepwise network construction procedures. Signed network construction considers only positively correlated nodes to be connected. Briefly, a correlation matrix of all genes and samples was created using Pearson’s correlation, which was then transformed to an adjacency matrix via soft-thresholding using the lowest power that maximizes scale-free topology fit. A topological overlap matrix (TOM) and topological overlap dissimilarity measure were then calculated to hierarchically cluster highly interconnected genes. These clusters were then partitioned into modules using the dynamic tree cut algorithm with a minimum module size of 30, and modules with similar expression profiles were merged when the module eigengene distance was above 0.25 (or correlation ≥ 0.75). Merged modules were then correlated with treatment, time, and infection load to identify network expression significantly associated with responses to Bd-Brazil and Bd-GPL infection over time.

Following pairwise differential expression and network co-expression analyses, functional enrichment tests were performed on all significant gene sets and modules using topGO (v 2.34.0). We restricted our analysis to only the Biological Process GO category, used all Trinotate-annotated genes as background, and performed all enrichment tests using the ‘weigh01’ algorithm and Fisher’s exact tests. We further examined host responses by focusing explicitly on genes involved in immunity. We filtered significantly expressed genes in all tests to only those with Blast annotations involved in innate immunity, adaptive immunity, and *a priori* amphibian chytrid responses using a custom R script.

## RESULTS

### Infection trial

Survival of frogs challenged with Bd-Brazil and Bd-GPL was 80% and 50%, respectively, which was marginally significant (p = 0.065) (Figure 1a). Bd-negative control frogs remained Bd-free throughout the experiment (Figure 1b). At early infection, neither treatment group varied significantly in infection intensity from controls (log10Bd- Brazil = 0; log10Bd-GPL = 2.09 ± 2.42, *t*[3] = 1.73, p = 0.18). At mid and late-stage infection, infection intensity varied significantly with treatment with Bd-GPL+ frogs harboring higher infection intensity [*F*[1,13] = 34.355, *p* < 0.0001]. Within-treatment, infection intensity did not vary with time between mid and late-stage infection [*F*[1,13] = 1.769, *p* = 0.206].

**Figure 1.**
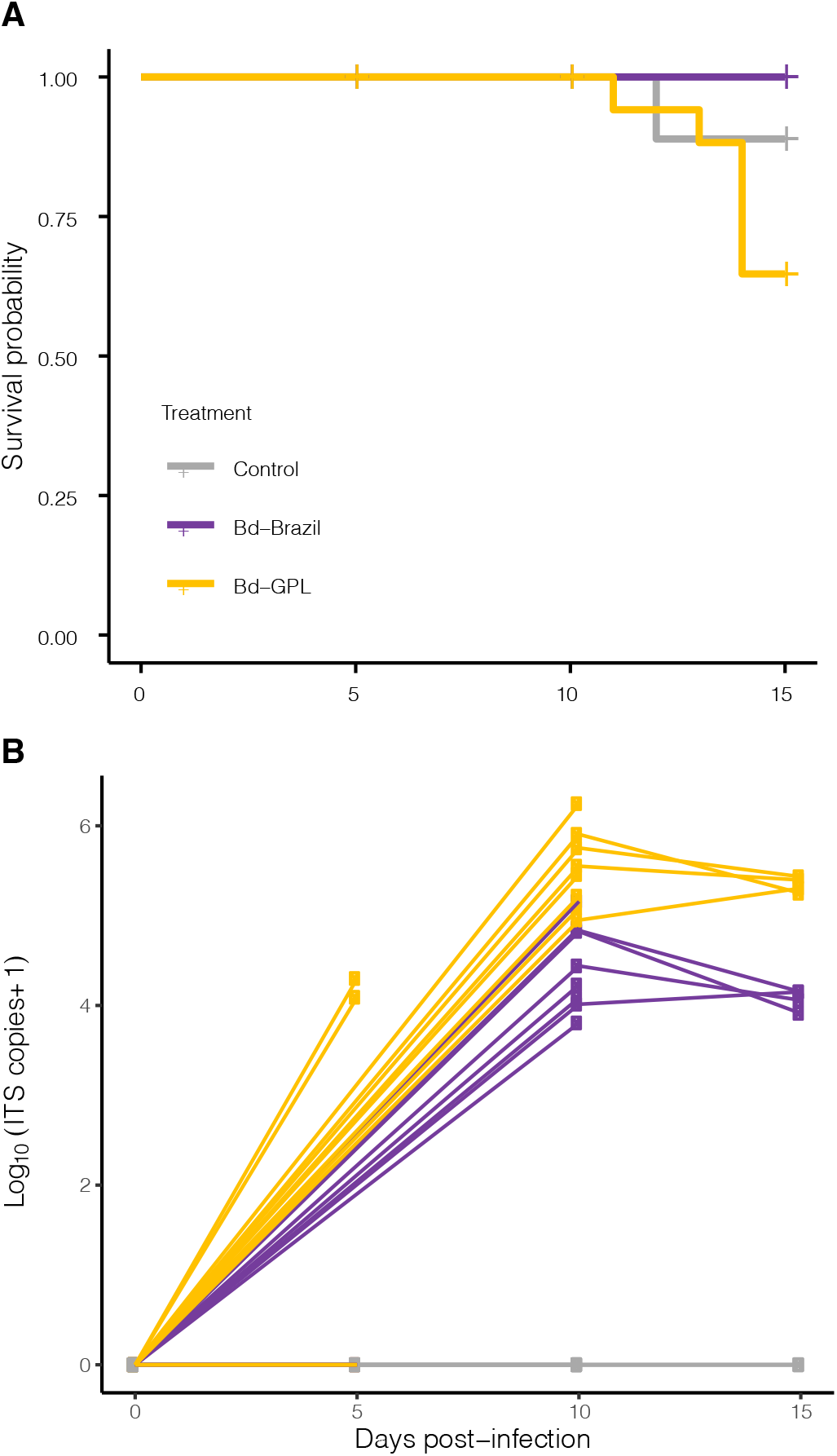
A) *Brachycephalus pitanga* survival probability over time, estimated using Kaplan-Meier method. B) Bd infection intensity over time, measured via number of ITS copies.

### Transcriptome assembly and annotation

Sequencing generated a total of 1.48 billion single-end and 844 million paired-end raw reads. Following filtering, we used 841 million paired-end reads from 36 samples for transcriptome assembly. Filtered single-end reads averaged 13.8 million reads (SD 3.2 million) per sample and were used for alignment and quantification. De novo Trinity assembly using samples from all tissues yielded a transcriptome of 298,388 Trinity genes. Our BUSCO analysis demonstrated high transcriptome completeness, with 91.1 % of conserved vertebrate orthologs observed (complete: 91.1% [single: 41.6%, duplicated: 49.4%], fragmented: 2.7%, missing: 6.2%, n = 3950). Using the Trinotate pipeline, we successfully annotated 73,386 (25%) Trinity transcripts with Blast homology and GO annotation. We removed 6,938 transcripts with non-vertebrate annotations and collapsed annotation to the gene level, which resulted in a cleaned, final transcriptome of 169,524 genes. We removed one liver and two intestinal samples that were clear outliers, retaining 105 total samples (Figure S1).

### Temporal variation in response

We analyzed differential expression in two ways: between treatment and control groups at each time point, and within treatment over time. In the latter analysis, we controlled for baseline temporal variation in gene expression by filtering out any genes that also varied significantly in control frogs over time. Overall, Bd-Brazil^+^ frogs mounted the largest change in expression response at mid-stage infection, while Bd-GPL^+^ frogs had the strongest response at late-stage infection (Figures 2a, 2b, S2). Compared to controls, Bd-Brazil^+^ frogs significantly differentially expressed 26 genes at early infection, 435 at mid-stage infection, and 9 at late-stage infection. In contrast, Bd-GPL^+^ frogs expressed 97 genes early, 86 genes at mid-stage infection, and 728 at late-stage infection. This expression response did not track with treatment-specific shifts in infection intensity; within each treatment, infection intensity did not vary significantly between mid- and late-stage infection (Figure 1). The trend of Bd-Brazil^+^ frogs expressing more genes at mid-infection and Bd-GPL^+^ frogs expressing more genes at late-stage infection was consistent both across all tissues and when considering immune-specific genes (Figures 2c, S3). Among genes with explicit roles in immunity, gene expression in treatment and control frogs varied most at mid- and late-stage infection (Figures 3b, 4b), with Bd-GPL^+^ frogs demonstrating the lowest expression correlation at late infection in skin (Figure S4).

**Figure 2.**
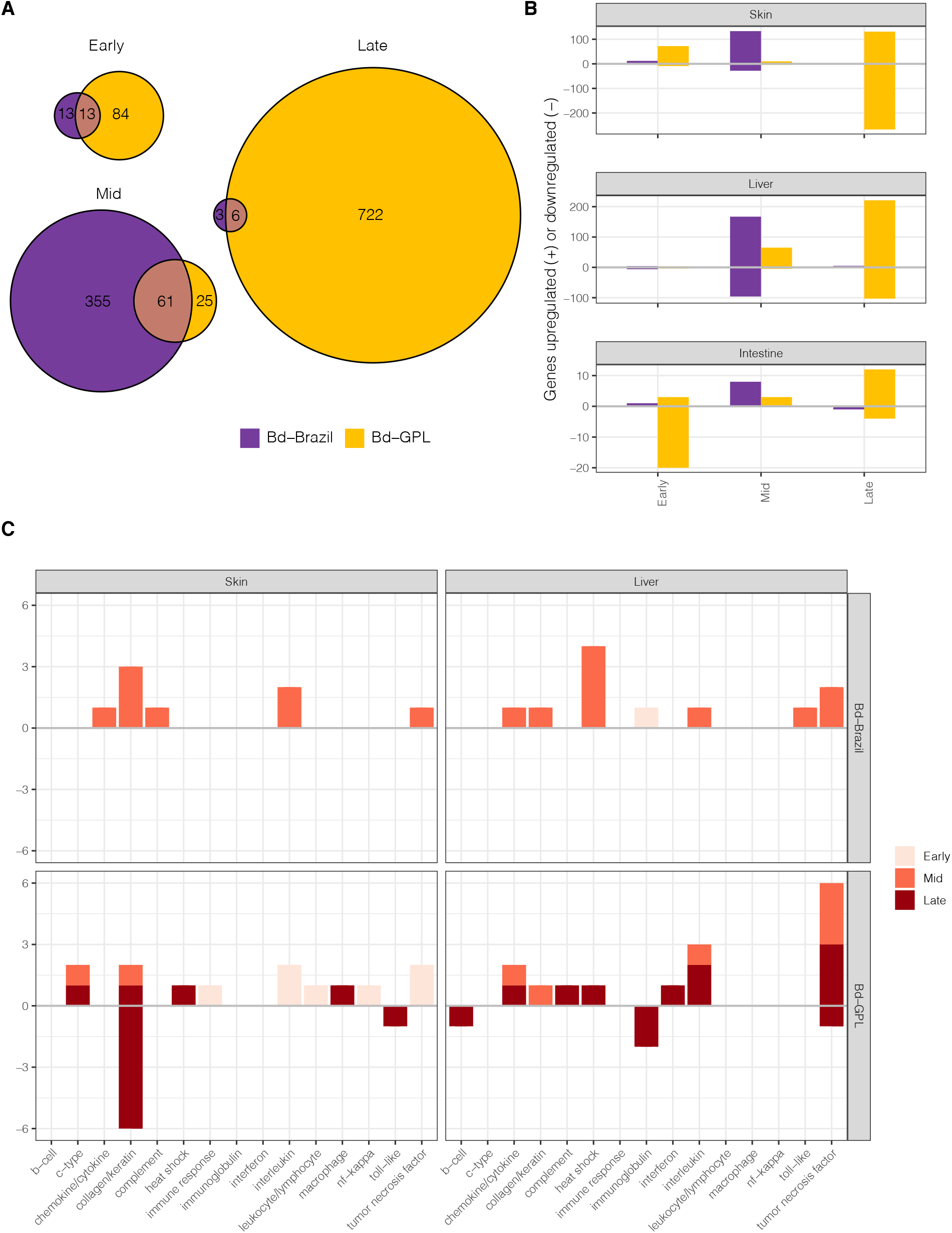
Bd-Brazil^+^ and Bd-GPL^+^ temporal variation in pumpkin toadlet gene expression. A) Area-proportional Euler plots showing number of genes differentially expressed relative to controls in each treatment over time. B) Number of genes differentially expressed relative to controls, separated by tissue. C) Number of immune genes differentially expressed over time, relative to controls.

**Figure 3.**
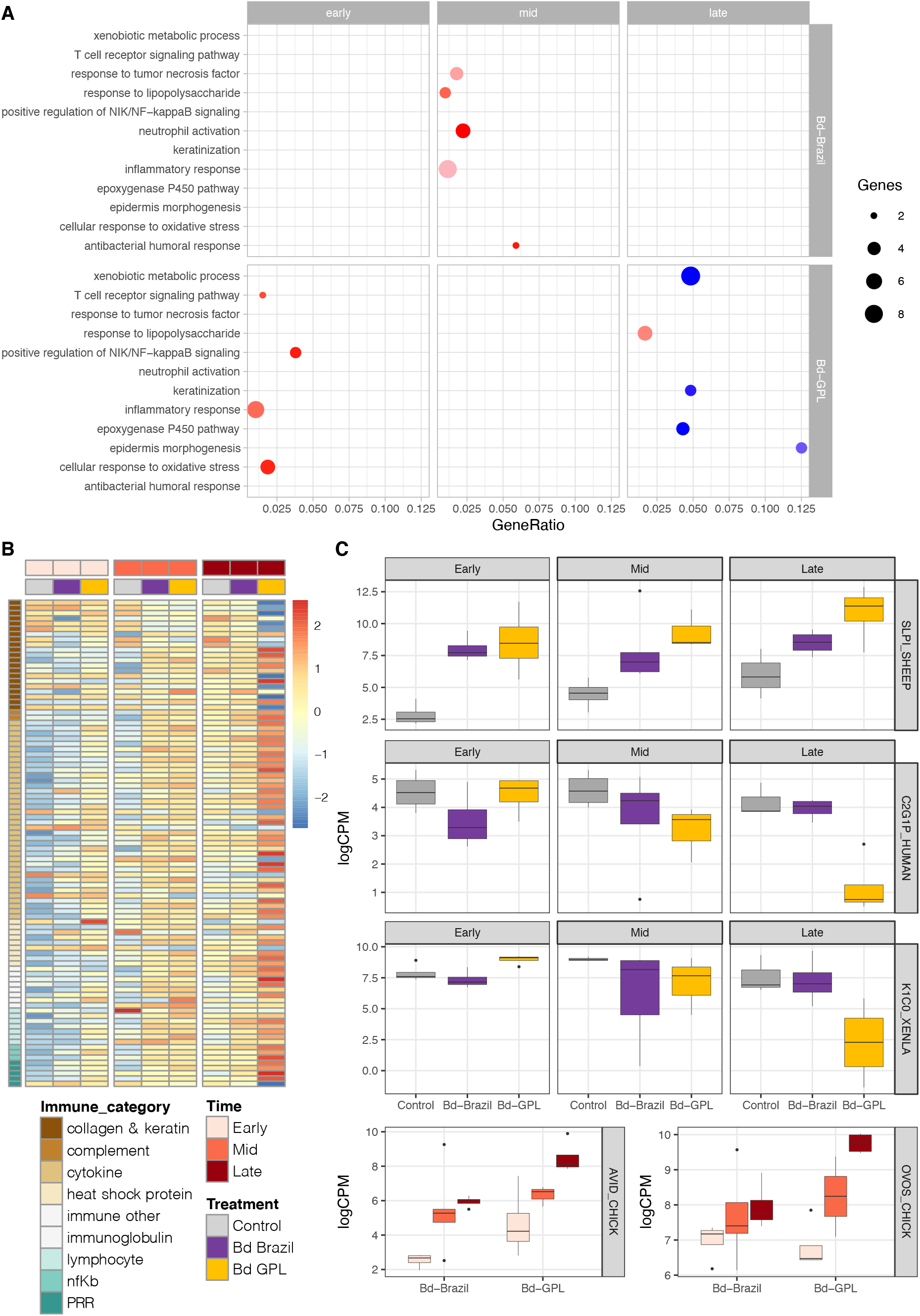
Differential responses to Bd-Brazil and Bd-GPL in the skin of *Brachycephalus pitanga*. A) Enrichment dotplot of selected GO terms related to immune response. Red represents upregulated terms, blue downregulated, and darker hues indicate higher significance. Full GO enrichment results can be found in File S2. B) Heatmap of immune genes over time. C) Selected immune genes uniquely significant to Bd-Brazil^+^ or Bd-GPL^+^ frogs at each time point. Included genes: SLPI (Antileukoproteinase), C2G1P (Putative inactive cytochrome P450 2G1), K1C0 (Keratin-3 type I cytoskeletal 51 kDa), AVID (Avidin), and OVOS (Ovostatin).

**Figure 4.**
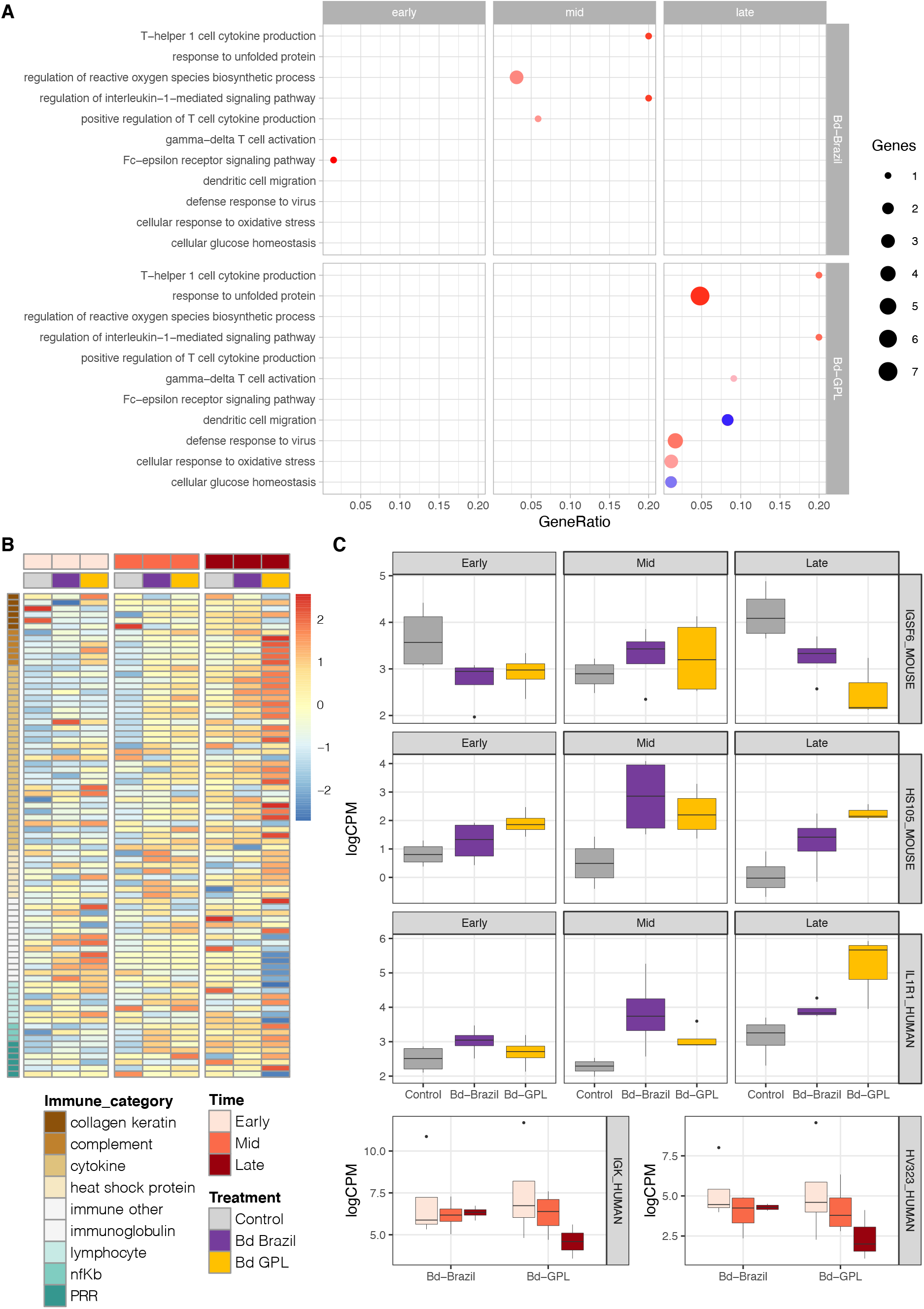
Differential responses to Bd-Brazil and Bd-GPL in the liver of *Brachycephalus pitanga*. A) Enrichment dotplot of selected GO terms related to immune response. Red represents upregulated terms, blue downregulated, and darker hues indicate higher significance. Full GO enrichment results can be found in File S2. B) Heatmap of immune genes over time. C) Selected immune genes uniquely significant to Bd-Brazil^+^ or Bd-GPL^+^ frogs at each time point. Included genes: IGSF6 (Immunoglobulin superfamily member 6), HS105 (Heat shock protein 105 kDa), IL1R1 (Interleukin-1 receptor type 1), IGK (Immunoglobulin kappa light chain), and HV323 (Immunoglobulin heavy variable 3-23).

### Shared responses to Bd

Despite key differences in timing and magnitude of host immune response to each lineage, Bd infection also activated core host responses regardless of lineage.

Comparatively few genes were expressed by both Bd-Brazil^+^ and Bd-GPL^+^ frogs at each stage; however, several of these were immune-related (Table 1). These generalized Bd responses may be important for the survival of *B. pitanga* in a region where multiple Bd lineages co-occur.

**Table 1.** Brachycephalus pitanga shared response to chytridiomycosis. Immune-related genes significantly expressed in both Bd-Brazil^+^ and Bd-GPL^+^ frogs at each time point.

| Trinity gene ID | Ortholog name | Tissue | Time | Bd-Brazil<br>logFC | Bd-GPL<br>logFC |
| --- | --- | --- | --- | --- | --- |
| TRINITY_DN535_c0_g1 | Antileukoproteinase | skin | early | 5.25 | 6.98 |
| TRINITY_DN5593_c0_g1 | Collagen alpha-1(XI) chain | skin | mid | 1.91 | 1.97 |
| TRINITY_DN1647_c0_g1 | Ovostatin | skin | mid | 2.32 | 2.70 |
| TRINITY_DN2885_c0_g1 | IgGfc-binding protein | skin | mid | 6.78 | 6.17 |
| TRINITY_DN623_c2_g2 | Solute carrier family 35 member E4 | skin | midvsearly | 1.51 | 1.83 |
| TRINITY_DN1354_c0_g2 | Heat shock 70 kDa protein | skin | midvsearly | 2.67 | 2.90 |
| TRINITY_DN21182_c0_g1 | ETS domain-containing protein Elk-3 | skin | latevsearly | 1.44 | 1.36 |
| TRINITY_DN13596_c0_g1 | CCN family member 1 | skin | latevsearly | 2.27 | 2.12 |
| TRINITY_DN130935_c0_g1 | Cartilage oligomeric matrix protein | skin | latevsearly | 3.13 | 2.16 |
| TRINITY_DN353_c0_g2 | Induced myeloid leukemia cell differentiation protein Mcl-1 homolog | liv | mid | 1.90 | 1.98 |
| TRINITY_DN3004_c0_g1 | Matrix metalloproteinase-14 | liv | mid | 2.14 | 2.20 |
| TRINITY_DN18284_c0_g2 | Tumor necrosis factor receptor superfamily member 12A | liv | mid | 2.44 | 2.68 |
| TRINITY_DN646_c0_g1 | Serine/threonine-protein kinase Sgk1 | liv | mid | 2.95 | 2.75 |
| TRINITY_DN1586_c0_g1 | Tumor necrosis factor receptor superfamily member 6B | liv | mid | 3.80 | 4.23 |
| TRINITY_DN4462_c0_g1 | Metalloproteinase inhibitor 1 | liv | mid | 4.14 | 4.63 |
| TRINITY_DN9842_c0_g1 | Transcription factor jun-B | liv | mid | 4.63 | 4.56 |
| TRINITY_DN10172_c0_g1 | Suppressor of cytokine signaling 3 | liv | mid | 5.77 | 5.58 |
| TRINITY_DN9170_c0_g1 | Metalloreductase STEAP1 | liv | late | 2.35 | 3.75 |
| TRINITY_DN14987_c0_g1 | Immunoglobulin heavy constant epsilon | liv | latevsearly | -7.22 | -6.68 |
| TRINITY_DN1636_c0_g1 | Interleukin-6 receptor subunit beta | liv | latevsearly | 1.56 | 2.16 |
| TRINITY_DN9170_c0_g1 | Metalloreductase STEAP1 | liv | latevsearly | 1.90 | 3.77 |
| TRINITY_DN12993_c0_g1 | Tumor necrosis factor receptor superfamily member 26 | liv | latevsearly | 2.90 | 3.12 |
| TRINITY_DN1586_c0_g1 | Tumor necrosis factor receptor superfamily member 6B | liv | latevsearly | 3.65 | 3.88 |

At early infection, Bd^+^ frogs upregulated very few shared genes, only one of which was involved in immunity. Antileukoproteinase, which protects epithelial tissues from serine proteases, was upregulated in skin (Figure 3c; SLPI_SHEEP) (Hiemstra et al., 1996). Although frogs upregulated two shared genes in liver, neither were annotated.

By mid-stage infection, Bd^+^ frogs upregulated ovostatin expression in skin, which also has serine-type endopeptidase inhibitor activity, as well as genes involved in metal ion binding, transmembrane transport, collagen, IgGFc-binding, and heat shock (Table 1). In liver, Bd^+^ frogs responded by upregulating multiple tumor necrosis factor receptors and a cytokine suppressor, as well as serine/threonine protein kinase, metalloproteinase inhibitor 1, and matrix metalloproteinase genes (Table 1).

When comparing late-stage responses to early responses in skin, Bd^+^ frogs continued to upregulate metalloproteinase inhibitor, cytokine signaling suppressor, and heat shock protein genes. Additionally, they upregulated genes involved in wound response (GO:0009611), cell proliferation (GO:0042127), and cell death (GO:0008219) (File S1). In liver, an immunoglobulin heavy chain gene was strongly downregulated, while tumor necrosis factor receptor, interleukin receptor, and metalloreductase genes were upregulated (Table 1).

Bd^+^ frogs exhibited no shared expression in intestine at early, mid, or late-stage infection.

### Unique responses to invasive and enzootic Bd

Although there were key shared responses to Bd infection, Bd-Brazil^+^ and Bd-GPL^+^ frogs expressed far more genes uniquely at each time point. Collectively, Bd-GPL^+^ frogs expressed more genes than Bd-Brazil^+^ frogs, indicating a stronger response to the invasive pathogen genotype (Figures 2a, 2b).

### Early infection

Bd-Brazil^+^ and Bd-GPL^+^ frog responses in skin at early infection were markedly different. Bd-Brazil^+^ frogs expressed very few genes, none of which could be annotated. In contrast, Bd-GPL^+^ frogs differentially expressed 70 genes, many of which were involved in both innate and adaptive immune responses. In particular, T cell receptor signaling pathway (GO:0050852), NF-kappaB regulation (GO:1901224), inflammatory response (GO:0006954), and cellular response to oxidative stress (GO:0034599) were among the significantly enriched biological processes upregulated by Bd-GPL^+^ frogs (Figure 3a). Genes explicitly related to immunity included an IgGFc-binding protein (TRINITY_DN2885_c0_g1, logFC = 5.70, FDR = 0.005), cornifelin homolog (TRINITY_DN270_c1_g1, logFC = 4.54, FDR = 0.003), matrix metalloproteinase-18 (TRINITY_DN8750_c0_g1, logFC = 4.02, FDR = 0.03), multiple tumor necrosis factor, NF-kB, and pro-inflammatory interleukin-17 genes (File S1), none of which were upregulated by Bd-Brazil^+^ frogs at this time.

Bd-Brazil^+^ and Bd-GPL^+^ frogs expressed few genes early in either liver or intestine. Among Bd-Brazil^+^ frogs, the only annotated gene was an Ig kappa chain, which was strongly upregulated in liver (TRINITY_DN76350_c0_g1, logFC = 6.22, FDR = 0.03). Bd-GPL^+^ frogs exhibited no significant expression change in liver, but downregulated three genes involved in immune response in intestine: a hepatitis A virus cellular receptor homolog involved in mast cell activation (TRINITY_DN5054_c0_g1, logFC = -5.91, FDR = 0.04), a heat shock protein (TRINITY_DN139560_c0_g1, logFC = -5.05, FDR=0.04), and a saxiphilin precursor (TRINITY_DN2018_c0_g1, logFC = -6.52, FDR = 0.03), which plays a putative role in microbial toxin neutralization (File S1) (Mahar et al., 1991).

### Mid Infection

By mid-stage infection, Bd-Brazil^+^ frogs ramped up responses in skin, while Bd-GPL^+^ frogs showed little response in skin at all. Bd-Brazil^+^ frogs upregulated 150 genes compared to control frogs and more than 500 genes within treatment compared to early infection. Relative to controls, genes were enriched for several immune processes including response to tumor necrosis factor (GO:0034612), response to lipopolysaccharide (GO:0032496), neutrophil activation (GO:0042119), inflammatory response (GO:0006954), and antibacterial humoral response (GO:0019731) (Figure 3a). Among the top 10 most upregulated genes compared to control frogs, 7 were involved in immunity (File S1). When comparing within treatment, GO terms such as epidermis development (GO:0008544) were significantly enriched, and interleukins, tumor necrosis factors, and other chemokines and cytokines were among the most upregulated genes at mid-stage infection relative to early infection (Files S1, File S2). In contrast, none of the immune genes that Bd-GPL^+^ frogs upregulated in skin at early infection were significantly expressed by mid-stage infection.

Bd-Brazil^+^ frogs also mounted a greater response than Bd-GPL^+^ frogs in liver. Among the 222 genes differentially expressed relative to controls and 35 within treatment between mid and early infection, many were involved in hepatic processes including cholesterol homeostasis (GO:006695) and lipid homeostasis (GO:0055088); the majority of these processes were downregulated (File S2). An Ig kappa chain ortholog (TRINITY_DN76350_c0_g1), which was upregulated in early infection, was downregulated by mid-stage infection when comparing within treatment (logFC = -5.31, FDR = 0.03). Simultaneously, relative to controls, Bd-Brazil^+^ frogs upregulated several immune-related genes including a toll-like receptor (TRINITY_DN2201_c0_g1, logFC = 7.17, FDR = 0.005), interleukin-1 receptor (TRINITY_DN8146_c0_g2, logFC = 1.89, FDR = 0.01), and multiple heat shock proteins and apoptotic proteins (Figure 4c, File S1). Much of this upregulation was enriched for immune processes such as T-helper 1 cell cytokine production (GO:0035744), regulation of reactive oxygen species biosynthetic process (GO:1903426), regulation of interleukin-1-mediated signaling (GO:2000659), and positive regulation of T cell cytokine production (GO:0002726) (Figure 4a).

Although Bd-GPL^+^ frog gene expression in liver was less pronounced with only 29 genes differentially expressed relative to controls and 93 genes between mid-stage and early infection, several genes were immune-related, including matrix metalloproteinase-18 (TRINITY_DN8750_c0_g1, logFC = 3.54, FDR = 0.04), a tumor necrosis factor receptor (TRINITY_DN12993_c0_g1, logFC = 2.3, FDR = 0.04), and an interleukin-3-regulated protein (TRINITY_DN24355_c1_g1, logFC = 2.08, FDR = 0.04). Within treatment, a class I histocompatibility antigen was one of the most downregulated genes by mid-stage infection (TRINITY_DN19868_c2_g1, logFC = - 3.62, FDR = 0.02) (File S1). Bd-GPL^+^ frog mid-stage gene expression relative to controls and within treatment was predominantly enriched for maintenance processes such as response to cAMP (GO:0051591) and intracellular signal transduction (GO:0035556) (File S2).

We found very little response in intestine at mid-stage infection in either treatment group. When comparing within treatment, Bd-Brazil^+^ frogs differentially expressed only 11 genes at mid-stage infection relative to early infection, one of which was a complement factor (TRINITY_DN2169_c0_g1, logFC = -3.73, FDR = 0.04). Bd- GPL^+^ frogs showed no differential expression either compared to controls or within treatment.

### Late Infection

By late-stage infection, Bd-Brazil^+^ frogs had ramped down immune responses in skin, while Bd-GPL^+^ frogs increased immune gene expression. Compared to control frogs, Bd-Brazil^+^ frogs showed no differential expression, while Bd-GPL^+^ frogs significantly expressed nearly 400 genes. Whereas the majority of genes differentially expressed at early and mid-stage infection were upregulated, by late-stage infection, Bd-GPL^+^ frogs downregulated more than 266 genes. These downregulated genes were enriched for keratinization (GO:0031424) and epidermis morphogenesis (GO:0048730); of the top 10 most downregulated genes, four were keratin orthologs (File S1). Downregulated genes were also enriched for xenobiotic metabolic process (GO: 0006805) and epoxygenase P450 pathway (GO:0019373), driven largely by decreased expression of several cytochrome P450 orthologs (Figure 3a, 3c). Upregulated genes were enriched for immune processes including response to lipopolysaccharide (GO:0032496) and included avidin (TRINITY_DN138572_c0_g1, logFC = 5.48, FDR=0.003), antileukoproteinase (TRINITY_DN535_c0_g1, logFC = 4.90, FDR = 0.04), and an IgGFc-binding protein (TRINITY_DN2885_c0_g1, logFC = 4.66, FDR = 0.04) among the most strongly upregulated genes (Figure 3a, 3c, File S1).

Within-treatment comparisons revealed similar responses in skin. Bd-Brazil^+^ frogs significantly differentially expressed 122 genes at late-stage infection compared to either early or mid-stage infection, while Bd-GPL^+^ frogs expressed more than 1,500 genes, the majority of which were downregulated. Relative to early infection, late-stage Bd-Brazil^+^ frogs upregulated genes involved in cell turnover (cell-matrix adhesion, GO:0007160; cell death, GO:0008219; cell motility, GO:0048870) (File S2). While Bd- GPL^+^ frogs demonstrated enrichment for similar cell migration and turnover processes, they also downregulated genes involved in epithelial maintenance. Specifically, genes with decreased expression were involved in keratinization (GO:0031424), cornification (GO:0070268), epidermal differentiation (GO:009913), and epithelial cell migration (GO:0010631). Of the 650 genes with increased expression by late-stage infection relative to early infection, there was significant enrichment for inflammatory response (GO:0006954), as well as multiple terms related to apoptosis, interleukin signaling and secretion, and tumor necrosis factor response and regulation (Files S1, S2).

This diminished response in Bd-Brazil^+^ and increased response in Bd-GPL^+^ frogs was mirrored in liver. By late-stage infection, Bd-Brazil+ frogs significantly expressed only three genes compared to control frogs, while Bd-GPL^+^ frogs expressed over 300.

One-third of these 300 genes were downregulated, and were involved in processes indicative of metabolic disruption, including multicellular organism development (GO:0007275) and cellular glucose homeostasis (GO:0001678). Of the two-thirds of genes that were upregulated, these were enriched for multiple immune-related processes that were enriched in Bd-Brazil^+^ frogs at mid-stage infection, such as T-helper 1 cell cytokine production (GO:0035744) and regulation of interleukin-1-mediated signaling pathway (GO:2000659). Bd-GPL^+^ frogs also upregulated genes enriched for defense response to virus (GO:0051607) and cellular response to oxidative stress (GO:0034599), as well as more general processes involved in cellular maintenance, such as response to unfolded protein (GO:0006986) (Figure 4a). Multiple tumor necrosis factor, interleukin, metalloproteinase, and complement factor genes were among those most upregulated (File S1).

When examining liver gene expression within treatment, Bd-Brazil^+^ frogs expressed far fewer genes than Bd-GPL^+^ frogs (66 versus 654 genes). Immunoglobulin gene expression was notably downregulated by Bd-GPL^+^ frogs. Whereas Bd-Brazil^+^ frogs decreased expression of one immunoglobulin gene by late-stage infection, four of the top 10 most downregulated genes by Bd-GPL^+^ were immunoglobulin orthologs, and B-cell antigen receptors and transcription factors were also downregulated (Figure 4c, File S1).

In intestine, Bd-Brazil^+^ frogs expressed two genes, while Bd-GPL^+^ frogs expressed 19, including three heat shock proteins, matrix metalloproteinase-18, and metalloproteinase inhibitor 1.

### 3.5.4 Late-stage Bd-GPL versus mid-stage Bd-Brazil

To examine whether Bd-GPL^+^ frog responses were similar to Bd-Brazil^+^ responses and only temporally shifted, we compared late-stage Bd-GPL^+^ responses directly to mid-stage Bd-Brazil^+^. Within skin, late-stage Bd-GPL^+^ frogs differentially expressed 139 genes compared to mid-stage Bd-Brazil^+^ frogs. As with comparisons of late-stage Bd- GPL^+^ frogs relative to controls, downregulated genes were enriched for processes including keratinization (GO:0031424), xenobiotic metabolic process (GO:0006805), and epoxygenase P450 pathway (GO0019373), with keratin and cytochrome P450 orthologs among the genes most downregulated by Bd-GPL^+^ frogs (File S1, S2). In liver, late-stage Bd-GPL^+^ frogs expressed 98 genes differentially, with downregulation of several zinc finger protein orthologs and upregulation of a cytokine signaling suppressor and C-X-C motif chemokine 14 (File S1, S2). No genes were significantly differentially expressed in intestine between the two treatment groups.

### Gene network analyses

Among WGCNA modules significantly correlated with treatment, time point comparison, and infection intensity in skin, nearly all modules were correlated with both treatment and at least one time point, indicating an interaction effect. Of these modules, we found three with extensive enrichment for immune responses. Only one module (MEgreen) was negatively correlated with treatment and not time, indicating increased expression in Bd-GPL^+^ relative to Bd-Brazil^+^ frogs at all time points. Other modules with increased expression in Bd-GPL^+^ frogs (MEblack, MEviolet) were also positively correlated with late and mid-stage infection relative to early infection (Figure 5b). Surprisingly, no modules were correlated with infection intensity (Figure S5).

**Figure 5.**
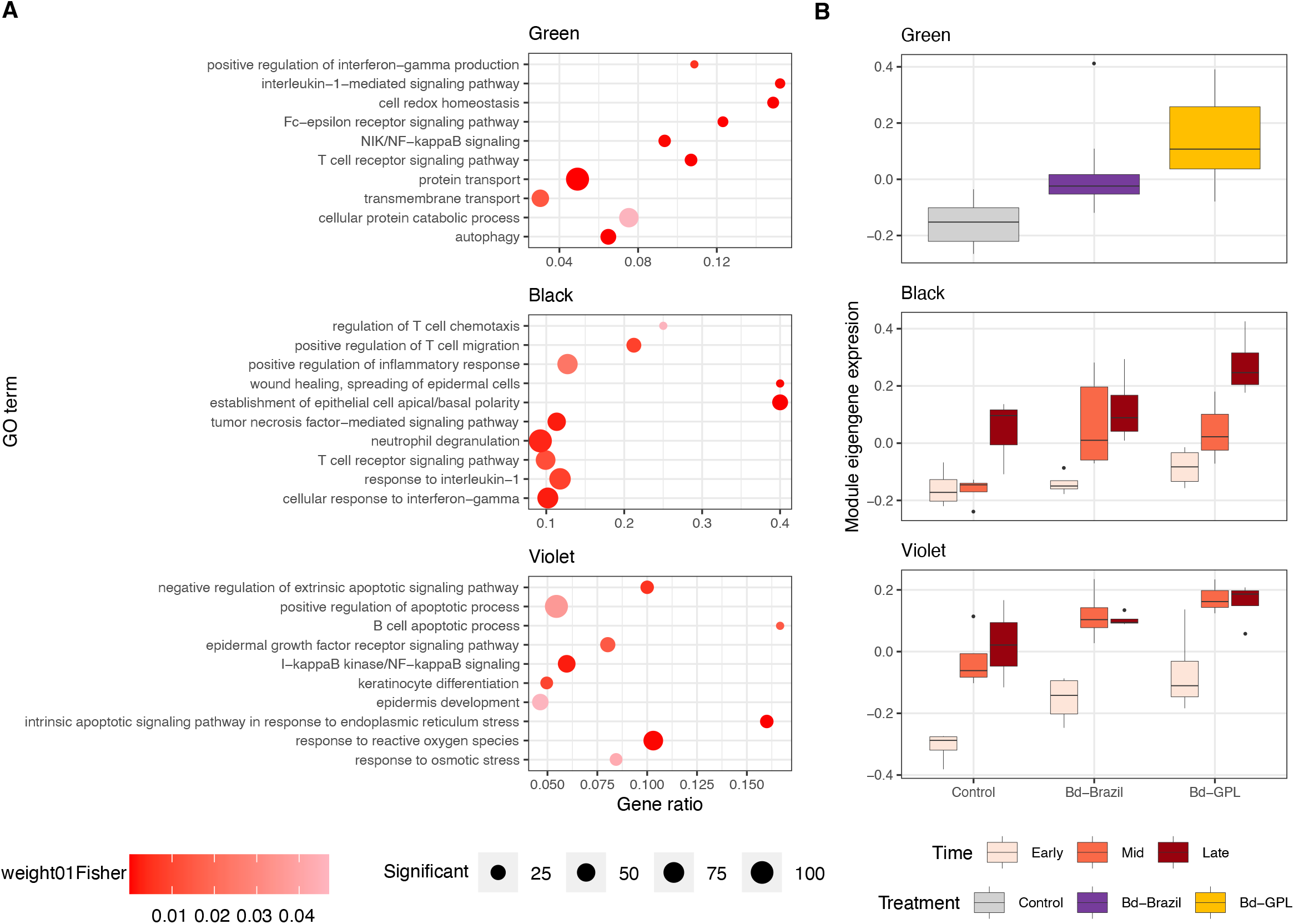
Gene network expression in the skin of *Brachycephalus pitanga*. A) Enrichment dotplot of selected GO terms significant in either the green, black, or violet WGCNA modules. Full GO enrichment results can be found in File S2. B) Module eigengene expression related to time and treatment. The green module was significantly correlated with treatment (Bd-GPL vs. Bd-Brazil), while the black and violet modules were correlated with treatment and time comparisons Early vs. Late and Early vs. Mid

All three skin modules showed broad enrichment for both innate and adaptive immune processes. Module MEgreen was a general immune response module, with significant enrichment for terms including Fc-epsilon receptor signaling pathway (GO:0038095), T cell receptor signaling pathway (GO:0050852), interleukin-1-mediated signaling pathway (GO:0070498), positive regulation of interferon-gamma (GO:0032729), and NIK/NF-kappaB signaling (GO:0038061), as well as for terms related to protein transport, folding, and catabolism (Figure 5a). In addition to several cytokine processes, module MEblack was enriched for inflammation, numerous T cell processes (e.g., T cell differentiation, GO:0030217), as well as epithelial cell development and differentiation (e.g. glandular epithelial cell development, GO:0002068 and wound healing, GO:0035313) (Figure 5a). Module MEviolet showed significant enrichment for multiple GO terms related to NF-kB signaling (e.g. I-kappaB kinase/NF-kappaB signaling, GO:0007249), pre-B cell differentiation (GO:0002329), and apoptosis (e.g. positive regulation of apoptotic process, GO:0043065) (Figure 5a).

Several modules were correlated with treatment, time comparison, and infection intensity in liver. Modules significantly correlated with infection intensity were all also significant for either treatment and/or time comparison, indicating that zoospore load, regardless of genotype, did not uniquely drive host expression networks (Figure 6b, S6). Among significantly correlated modules, only two showed widespread enrichment for immune responses. Module MEsaddlebrown, which was negatively correlated with treatment and positively correlated with each time comparison, showed enrichment broadly for cell division (GO:0051301) and migration (GO:0030335), and specifically for T cell regulation (GO:0046007) and B cell activation (GO:0042113) (Figure 6a). Module MEcyan was positively correlated with treatment and negatively correlated with early versus mid and late versus mid time comparisons. This module was more generally involved in immune response, with enrichment for terms including adaptive immune response (GO:0002250), complement activation (GO:0006958), neutrophil degranulation (GO:0043312), and Fc-gamma (GO:0038096) receptor signaling (Figure 6a).

**Figure 6.**
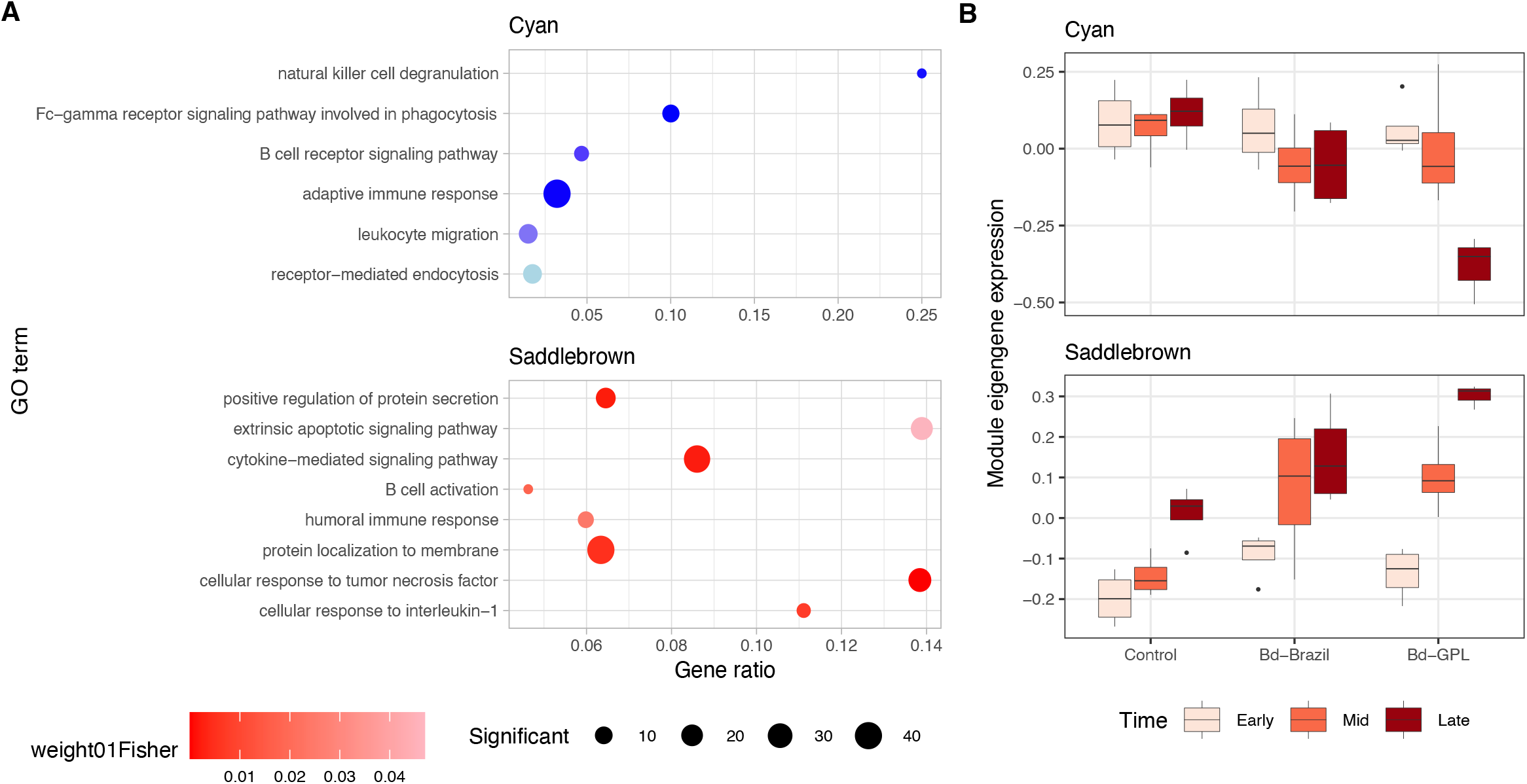
Gene network expression in the liver of *Brachycephalus pitanga*. A) Enrichment dotplot of selected GO terms significant in either the green, black, or violet WGCNA modules. Full GO enrichment results can be found in File S2. B) Module eigengene expression related to time and treatment. The green module was significantly correlated with treatment (Bd-GPL vs. Bd-Brazil), while the black and violet modules were correlated with treatment and time comparisons Early vs. Late and Early vs. Mid.

## DISCUSSION

Our results show that in the endemic, direct-developing pumpkin toadlet, *Brachycephalus pitanga*, host immune response strategies varied with exposure to divergent pathogen lineages. Hosts were less susceptible to enzootic Bd-Brazil, mounting a successful immune response by mid-stage infection that was dominated by epithelial turnover and innate immune activation. This response was fully resolved by late-stage disease, even though Bd-Brazil infection load remained constant.

Conversely, when frogs were exposed to the highly virulent Bd-GPL, responses were both magnified and delayed, with largescale downregulation of keratin, cytochrome P450, and immunoglobulin genes at late-stage infection. Thus, whereas previous studies have shown variation in immune responses between susceptible versus resistant species, infected versus control frogs, and in susceptible hosts upon re-exposure, our results highlight that within the same species, host immune responses can shift profoundly depending on Bd lineage.

In the past several decades, animal infectious disease biologists have increasingly recognized the importance of decoupling host susceptibility to parasitism into measures of resistance and tolerance (Best et al., 2014). Resistance, or the ability of a host to avoid infection or decrease parasite density once infected, has different effects on epidemiological and coevolutionary dynamics than tolerance, defined as the host’s ability to limit negative consequences of infection without affecting parasite density (Boots et al., 2009; Råberg et al., 2007, 2009b). The former is predicted to result in antagonistic coevolution between host and parasite, whereas the latter is predicted to not constrain parasite fitness, instead allowing for increased parasite prevalence, leading to selection for and potentially eventual fixation of tolerance alleles (Roy & Kirchner, 2000).

Whereas resistance can lead to classical antagonistic coevolution between pathogen and host, tolerance coevolution can result in multiple stable evolutionary states (Best et al., 2014). In particular, while tolerance may allow for apparent commensalism wherein both host and pathogen minimize virulence, it can also lead to evolution of parasites that are highly virulent toward nontolerant hosts, resulting in catastrophic outbreaks (Miller, 2006). Understanding the mechanisms of host susceptibility, tolerance, and resistance is therefore key to characterizing not only individual disease trajectories, but population level outcomes.

In this study, host survival was higher for Bd-Brazil^+^ than Bd-GPL^+^ frogs. Zoospore loads were significantly lower in Bd-Brazil^+^ than Bd-GPL^+^ frogs at each time point but did not vary within treatment from mid- to late-stage infection. Differential survival rates were marginally significant; increased survival combined with decreased zoospore load suggests a degree of resistance and/or tolerance of pumpkin toadlets to enzootic Bd. That zoospore load did not decrease between mid and late-stage infection could suggest tolerance, although the short duration of our trial does not allow us to determine whether toadlets are able to reduce infection loads over longer time periods. Because Bd-Brazil is enzootic to the pumpkin toadlet geographic range, yet periodic Bd- related host population declines have not been observed in this species, it is possible that the two species have coevolved under a tolerance framework, achieving a stable equilibrium wherein lower virulence Bd-Brazil persists. This could further explain why Bd-Brazil is outcompeted by Bd-GPL, yet maintains high virulence in a non-native host (Becker et al., 2017; Jenkinson et al., 2016, 2018). Further studies of Bd-Brazil prevalence in wild populations and longer-term infection dynamics in laboratory trials will help us delineate the extent to which resistance and/or tolerance account for pumpkin toadlet persistence.

Although overall patterns of immune activation diverged with pathogen lineage, frogs also shared key anti-Bd responses throughout infection. Early upregulation of antileukoproteinase in skin, the site of infection, may be especially pertinent during Bd infection. Antileukoproteinase is a small protein produced by cells at the mucosal lining that has both antimicrobial and antifungal properties. It acts locally at the mucosal epithelium to limit damage from inflammation by inhibiting proteolytic enzyme activity, and has particular efficacy against serine proteases (Vandooren et al., 2018). Serine-type peptidases are notably expanded in the Bd genome, are expressed by both Bd- Brazil and Bd-GPL lineages, and may be key to the transition to pathogenicity within the *Batrachochytrium* genus (Joneson et al., 2011; McDonald et al., 2019). Shared early upregulation of a serine protease inhibitor by frogs could indicate a general response combatting nonspecific Bd colonization of the skin epithelium.

General shared responses by mid- and late-stage infection were centered on serine protease inhibition, inflammation, and epithelial maintenance in skin, and increased cytokine and matrix metalloproteinase production in liver. IgGFc binding protein, which is involved in mucosal maintenance, was strongly upregulated in skin at mid-stage infection. At both mid- and late-stage infection, tumor necrosis factor receptor subfamily 6b, also known as decoy receptor 3 (DcR3), was upregulated in liver. This receptor is a biomarker of acute and chronic inflammation, and a therapeutic target for attenuating a number of autoimmune diseases in humans (Lin & Hsieh, 2011).

Whereas the majority of shared immune genes were upregulated, by late-stage infection immunoglobulin heavy constant epsilon, a key antibody component, was strongly downregulated in both Bd-Brazil^+^ and Bd-GPL^+^ frogs. Collectively, shared gene expression in both treatments suggests consistent inflammatory and epithelial damage responses, with concurrent late-stage humoral downregulation.

Lineage-specific responses to Bd-Brazil and Bd-GPL varied in timing, magnitude, and direction of expression. Unlike in Bd-Brazil^+^ frogs, wherein immune responses were generally upregulated, Bd-GPL^+^ frogs downregulated a multitude of genes. Epidermal cell differentiation, wound healing, and xenobiotic metabolic processes were among the most significantly downregulated gene ontology pathways. Keratin downregulation in skin has been observed previously in comparisons of susceptible versus resistant host species (Ellison, Savage, et al., 2014; Poorten & Rosenblum, 2016; Rosenblum, Poorten, Settles, et al., 2012). Bd colonization is limited to the skin, which leads clinically to hyperkeratosis, hyperplasia, and increased skin sloughing, in turn compromising requisite epithelial function in osmotic exchange and ion balance (Berger et al., 2005; Voyles et al., 2009). Similar to other susceptible hosts, we found that Bd- GPL^+^ frogs downregulated keratin gene expression at late infection. In contrast, Bd- Brazil^+^ frogs increased epidermal maintenance at mid-stage infection, which they then resolved by late infection. This resolution suggests a degree of Bd-Brazil tolerance because there was no concurrent change in infection intensity.

Bd-GPL^+^ frogs also downregulated several cytochrome P450 orthologs in skin. Whereas cytochrome P450 enzymes are predominantly found in liver, they are also active in skin, where they perform the same function in oxidative xenobiotic clearance (Baron et al., 2008). The fact that we only observed P450 downregulation in skin is expected given that this is the site of Bd infection. Bd is known to secrete a number of proteases and lymphotoxic virulence factors (Farrer et al., 2017; Fites et al., 2013; Rosenblum, Poorten, Joneson, et al., 2012). Downregulation of cytochrome P450 enzyme expression in the face of Bd-GPL infection suggests that frogs may be less able to clear these compounds, potentially leading to clinical disease and mortality.

By late-stage infection, when lineage-specific differences in mortality were apparent, Bd-GPL^+^ frogs mounted extensive hepatic expression responses. These results agree with previous histopathological findings in Bd-susceptible species, wherein infection led to leukocyte infiltration and hepatic atrophy (Salla et al. 2020). Most notably, whereas upregulated cytokine responses were similar to those observed at mid-stage Bd-Brazil infection, late-stage responses to Bd-GPL involved marked downregulation in a number of adaptive immune response genes. These results are consistent with previous studies that have found late-stage lymphocyte suppression in a number of susceptible populations and species, further demonstrating that Bd-mediated immunosuppression may dictate pathogenesis (Ellison, Savage, et al., 2014; Ellison, Tunstall, et al., 2014; Grogan, Cashins, et al., 2018).

Our results of late-stage immune gene expression variation in Bd-GPL^+^ frogs mirror those found in previous studies of susceptible frog species. Among studies that measured infection over time, Grogan, Cashins, et al. (2018) found that transcriptional responses in *Litoria verreaxii* were most up or downregulated in skin at their latest sampling point (14 days post-infection), and that the number of significantly expressed genes was higher in more susceptible populations. Skin gene expression in the endangered *Rana muscosa* and *Rana sierrae* also increases in late-stage relative to early disease (Rosenblum, Poorten, Settles, et al., 2012). Profound increases in the number of genes expressed at late-stage infection in susceptible relative to resistant species have also been observed in other studies that measured expression at a single, end-stage time point (Ellison, Tunstall, et al., 2014; Poorten & Rosenblum, 2016).

Both in skin and systemically, late-stage immune responses appear nonprotective in multiple susceptible species, and thus are considered dysregulated. The degree to which this dysregulation is pathogen-mediated, a function of immunopathology, or some combination of the two is an open question. Our results are consistent with previous studies that have documented some level of Bd-mediated adaptive immunosuppression in susceptible species at late-stage disease (Ellison, Savage, et al., 2014; Ellison, Tunstall, et al., 2014; Grogan, Cashins, et al., 2018).

Whereas we did not sample spleen and therefore predictably did not identify extensive systemic T cell expression, we did find downregulation of multiple immunoglobulin orthologs in liver. Although Bd is intracellular and thus adaptive responses are thought to be largely cell-mediated, shifts in humoral gene expression have been previously documented as well (Ellison, Savage, et al., 2014; Ellison, Tunstall, et al., 2014; Poorten & Rosenblum, 2016). Whereas Ellison et al. (2014) documented immunoglobulin upregulation in both skin and spleen of *Atelopus zeteki* upon re-exposure to Bd, Poorten et al. (2016) showed that susceptible *Anaxyrus boreas* decreased immunoglobulin expression in liver. Our results highlight extensive host immunoglobulin downregulation upon exposure to virulent Bd-GPL, but little to no downregulation when exposed to enzootic Bd-Brazil.

As opposed to adaptive immunosuppression in susceptible hosts, we found limited evidence of immunopathology. Although the absolute number of genes expressed was greatest in Bd-GPL^+^ frogs at late-stage infection, the degree to which these genes are involved in immunopathology is unclear. Specifically, we found similar numbers of pro-inflammatory cytokines significantly upregulated by Bd-Brazil^+^ and Bd- GPL^+^ frogs over the course of infection, although Bd-Brazil+ frogs typically expressed them earlier. Bd-Brazil^+^ frogs did upregulate expression of more cytokine suppressor 3 orthologs in both skin and liver than did Bd-GPL^+^ frogs. Thus, whereas pro-inflammatory cytokine expression was similar in gene number, suppression of cytokine signaling was more robust in Bd-Brazil+ frogs, perhaps suggesting increased mediation of inflammatory responses. However, it is unclear whether comparatively minimal cytokine suppressor 3 expression in Bd-GPL^+^ frogs is pathogen-mediated or a hallmark of immunopathology. GO term enrichment related to inflammation was similar between Bd-Brazil^+^ and Bd-GPL^+^ frogs, again calling immunopathology as a mechanism of pathogenesis into question.

As with studies that profiled immune responses in Bd-resistant species, we found less immune activation in our lower virulence, Bd-Brazil^+^ treatment. Because most gene expression studies include only end-stage tissue sampling, it is possible that resistant species have similarly timed immune responses to those we report in Bd-Brazil^+^ frogs, and that immunomodulation occurs earlier than was captured by sampling. Alternatively, the trajectories of low-susceptibility and resistant immune responses could be divergent. The only study with matched sampling of resistant and susceptible frogs over time found evidence of immune activation early in resistant frogs, but as with susceptible frogs, the magnitude of immune response was greatest at late-stage infection (Grogan, Cashins, et al., 2018). Thus, the trajectory of Bd-Brazil+ frogs, with immune activation occurring predominantly at mid-infection and nearly completely resolved by late-stage infection, indicates that temporal responses among resistant and low-susceptibility hosts differ.

Our results help clarify chytridiomycosis outcomes by explicitly examining lineage-specific responses in an endemic host species native to the sympatric range of enzootic Bd-Brazil and invasive Bd-GPL. Mechanisms of host Bd-Brazil susceptibility are marked by localized epithelial turnover and local and systemic innate immune activation that is largely resolved by late-stage infection, even in the face of stable zoospore infection intensities. Meanwhile, infection with virulent Bd-GPL elicits a non-protective response that is both delayed and more severe in magnitude, with widespread downregulation of keratin, cytochrome P450, and immunoglobulin genes. Thus, it is not only the specific immune pathways activated, but the timing of immune activation that determines disease outcome. In the context of host-pathogen coevolution, these mechanisms of Bd-Brazil susceptibility may be both key to host persistence in Brazil’s Southern Atlantic Forest and may have driven Bd-Brazil evolution toward a lower virulence optimum. As global spread of the hypervirulent Bd-GPL lineage has led to competition between Bd lineages, this lower virulence of Bd-Brazil could be linked to its competitive exclusion, further complicating epidemiology in the region.

## FUNDING

Grants and fellowships were provided by São Paulo Research Foundation (FAPESP #2016/25358-3; #2019/18335-5), the National Council for Scientific and Technological Development (CNPq #300896/2016-6; #302834/2020-6), the Coordination for the Improvement of Higher Education Personnel (CAPES - Finance Code 001), and a Cornell University Graduate Research Travel Grant.

## ACKNOWLEDGEMENTS

The authors thank Miranda M. Gray for invaluable assistance with experimental design and logistics, Andréa F.C. Mesquita for laboratory assistance, Cinnamon S. Mittan, Jordan Garcia, Maria Akopyan, David A. Chang van Oordt, Anat M. Belasen, and Lina M. Arcila Hernandez for manuscript feedback, and the Haddad lab for field assistance.

## Notes

### Competing Interest Statement

The authors have declared no competing interest.

